# Extremely rare variants reveal patterns of germline mutation rate heterogeneity in humans

**DOI:** 10.1101/108290

**Authors:** Jedidiah Carlson, Adam E Locke, Matthew Flickinger, Matthew Zawistowski, Shawn Levy, The BRIDGES Consortium, Richard M Myers, Michael Boehnke, Hyun Min Kang, Laura J Scott, Jun Z Li, Sebastian Zöllner

## Abstract

A detailed understanding of the genome-wide variability of single-nucleotide germline mutation rates is essential to studying human genome evolution. Here we use ∼36 million singleton variants from 3,560 whole-genome sequences to infer fine-scale patterns of mutation rate heterogeneity. Mutability is jointly affected by adjacent nucleotide context and diverse genomic features of the surrounding region, including histone modifications, replication timing, and recombination rate, sometimes suggesting specific mutagenic mechanisms. Remarkably, GC content, DNase hypersensitivity, CpG islands, and H3K36 trimethylation are associated with both increased and decreased mutation rates depending on nucleotide context. We validate these estimated effects in an independent dataset of ∼46,000 *de novo* mutations, and confirm our estimates are more accurate than previously published estimates based on ancestrally older variants without considering genomic features. Our results thus provide the most refined portrait to date of the factors contributing to genome-wide variability of the human germline mutation rate.

---

Germline mutagenesis is a fundamental biological process, and a major source of all heritable genetic variation (see Segurel et al.^1^ for a review). Mutation rate estimates are widely used in genomics research to calibrate variant calling algorithms^2^, infer demographic history^3^, identify recent patterns of genome evolution^4^, and interpret clinical sequencing data to prioritize likely pathogenic mutations^5^. Although mutation is an inherently stochastic process, the distribution of mutations in the human genome is not uniform and is correlated with genomic and epigenomic features including local sequence context^6, 7^, recombination rate^8^, and replication timing^9^. Hence, there is considerable interest in studying the regional variation and context dependency of mutation rates to understand the basic biology of mutational processes and to build accurate predictive models of this variability.

The gold standard for studying the germline mutation rate in humans is direct observation of *de novo* mutations from family-based whole-genome sequencing (WGS) data^9–12^. These studies have produced accurate estimates of the genome-wide average mutation rate (∼1 – 1.5 × 10^−8^ mutations per base pair per generation), and uncovered the aforementioned mutagenic effects of genomic features. However, given the inherently low germline mutation rate, family-based WGS studies detect only 40-80 *de novo* mutations for each trio sequenced^9,10,12^. Due to the sparsity of these observed mutations, it is difficult to accumulate a large dataset to precisely estimate mutation rates and spectrum at a fine scale and identify factors that explain genome-wide variability in mutation rates.

Other data sources for studying mutation patterns include between-species substitutions or within-species polymorphisms^7,8,13–16^. However, because these variants arose hundreds or thousands of generations ago, their distribution patterns along the genome have been influenced by the subsequent long-term actions of many evolutionary forces, such as natural selection and GC-biased gene conversion (gBGC), a process in which recombination-induced mismatches are preferentially repaired to G/C base pairs, resulting in an overabundance of common A/T-to-G/C variants^11,17,18^. To minimize the confounding effects of selection, studies that estimated mutation rates from these data tended to focus on intergenic non-coding regions of the genome, which are less often the target of selective pressure. Nevertheless, even putatively neutral loci may be under some degree of selection^19–21^, and are susceptible to the confounding effects of gBGC. Consequently, these processes bias the resulting distribution of variation, making it difficult to determine which trends are attributable to the initial mutation processes, and which to subsequent evolutionary factors. A further complication of estimating mutation rates with common variants is that the endogenous mutation mechanisms themselves have likely evolved over time^22^, so patterns of variation observed among these data may not necessarily reflect the same processes that are ongoing in the present-day population.

We therefore adopt an approach that relies exclusively on extremely rare variants (ERVs) to study innate mutation patterns across the genome. Here we exploit a collection of ∼35.6 million singleton variants discovered in 3,560 sequenced individuals from the BRIDGES study of bipolar disorder (corresponding to a minor allele frequency of 1/7120=0.0001404 in our sample). Compared to between-species substitutions or common variants in humans, these ERVs are extremely young on the evolutionary timescale (for a comparably-sized European sample, Fu et al. (2012) estimated the expected age of a singleton to be 1,244 years^23^), making them much less likely to be affected by evolutionary processes other than random genetic drift^1,11,17,24^. ERVs thus represent a relatively unbiased sample of recent mutations and are far more numerous than *de novo* mutations collected in family-based WGS studies.

Our results show that mutation rate heterogeneity is primarily dependent on the sequence context of adjacent nucleotides, confirming the findings of previous studies^7,9,25^. However, we demonstrate that our ERV-derived mutation rate estimates can differ substantially from estimates based on ancestrally older variants. Evaluating these differences in an independent dataset of ∼46,000 *de novo* mutations, collected from two published family-based WGS studies^9, 12^, we find that ERV-derived estimates yield a significantly more accurate portrait of present-day germline mutation rate heterogeneity. We further refine these estimates of context-dependent mutability by systematically estimating how mutation rates of different sequence motifs may be influenced by genomic features in wider surrounding regions, including replication timing, recombination rate, and histone modifications. Remarkably, we find that the direction of effect for certain genomic features often depends on the actual sequence motif surrounding the mutated site, underscoring the importance of jointly analyzing sequence context and genomic features. Accounting for these granular effects of the genomic landscape provides even greater accuracy in describing patterns of variation among true *de novo* mutations. Our results suggest that trends of variation throughout the genome are shaped by a diverse array of context-dependent mutation pathways, many of which have yet to be fully characterized. This high-resolution map of mutation rate estimates, along with estimates of the mutagenic effects of genomic features, is available to the community as a resource to facilitate further study of germline mutation rate heterogeneity and its implications for genetic evolution and disease.

## Results

### ERV data source and quality control

In the *Bipolar Research in Deep Genome and Epigenome Sequencing (BRIDGES)* study, we sequenced the genomes of 3,716 unrelated individuals of European ancestry to an average diploidgenome coverage of 9.6x (**Methods**). We identified and removed 156 samples which appeared to be technical outliers, resulting in a final call set of 35,574,417 autosomal ERVs from 3560 individuals (**Methods**). Due to the relatively low coverage of our sample, we likely failed to detect millions more ERVs—a recent study^26^ estimated the discovery rate for singletons in a sample of 4,000 whole genomes at 10x coverage to be ∼65-85%. Quality control measures indicate that the ERVs we detected are high quality, with a Transition/Transversion (Ts/Tv) ratio of 2.00, within the commonly observed range for single nucleotide variants (SNVs) from WGS data^27^ (**Supplementary Table 1**). Application of the 1000G strict accessibility mask^28^ (which delineates the most uniquely mappable regions of the genome) or a more stringent mapping quality score filter (MQ>56) did not appreciably change the Ts/Tv ratio (1.97-2.01) (**Supplementary Table 1**). We estimate fewer than 3% of the 35,574,417 ERVs are false positives (**Supplementary Note**), similar to the validated singleton error rates of other sequencing studies using a similar technology^28–30^. In addition, we present evidence that erroneous calls among the ERVs are unlikely to be biased by motif-specific genotyping error, mapping error, or mispolarization (**Supplementary Note**).

### Context-dependent variability in mutation rates

Prior studies have found that the nucleotides surrounding a mutated site are an important predictor of variability in mutation rates across the genome^7,11,25^. The most detailed such analysis to date, by Aggarwala and Voight^7^, considered the nucleotides up to 3 positions upstream and downstream from a variant site (i.e., a 7-mer sequence context), and estimated substitution probabilities per heptameric motif using 7,051,667 intergenic SNVs observed in 379 Europeans from phase 1 of the 1000 Genomes Project (hereafter referred to as the “1000G mutation rate estimates”). These estimates, though demonstrably more refined than mutation rates estimated in a 3-mer or 5-mer sequence context, have the potential problem of being derived from variants across the entire frequency spectrum. Among the 1000G SNVs used to estimate these rates, singletons and doubletons account for only ∼25%^7^, while the majority of variants occur at a higher frequency and thus likely arose hundreds or thousands of generations in the past. Over such a long time span, variants affected by cryptic selection, gBGC, or other evolutionary processes are more likely to have been fixed or disappeared, altering the distribution of observable variation.

Because ERVs are assumed to have occurred very recently in human history, we asked if ERV-based mutation rate estimates differed from the 1000G estimates, and if so, whether our revised estimation strategy would lead to more accurate representation of the basal mutation processes. To answer these questions, we first used the BRIDGES ERVs to estimate mutation rates according to mutation type (e.g., A>C, A>G, and so on) and local sequence context, considering the bases up to 3 positions upstream and downstream from each variant site (**Methods**). We refer to a mutation of a given type centered at a given sequence motif as a “mutation subtype” (e.g., C[A>C]G is a 3-mer subtype). Note that we are not estimating an absolute per-site, per generation mutation rate, but rather the relative fraction of each subtype containing an ERV within the BRIDGES data. We refer to rates calculated in this manner as “relative mutation rates,” and estimated these rates for all possible 1-, 3-, 5-, or 7-mer subtypes (**Supplementary Tables 2a-2d**).

ERV-derived relative mutation rate estimates for the six basic 1-mer mutation types (**Supplementary Table 2a**) reflect the expected higher mutability for transitions (A>G and C>T) relative to transversions (A>C, A>T, C>A, and C>G types)^1^. Splitting each mutation type into more granular subtypes reveals how additional patterns of mutation rate heterogeneity emerge as broader sequence contexts are incorporated (**Fig. 1; Supplementary Fig. 1**). Our ERV-based relative mutation rate estimates confirm nearly all of the hypo- or hypermutable motifs previously reported by Aggarwala and Voight^7^ and Panchin et al.^13^. A subset of these are highlighted in **Fig. 1a**, including lower relative mutation rates for NNN[C>T]GCG subtypes and A>G subtypes in motifs containing runs of 4 or more A bases (shown in green boxes), and higher relative mutation rates for N[A>G]T, N[C>T]G, and CA[A>G]TN subtypes (pink boxes). A particularly notable example of context-dependent hypermutability is the set of NTT[A>T]AAA subtypes (**Fig. 1b**), also described previously^7^. Despite A>T mutations having the lowest relative mutation rate among 1-mer types, its NTT[A>T]AAA subtypes have a >6-fold higher rate than the 1-mer A>T relative mutation rate.

**Figure 1.**
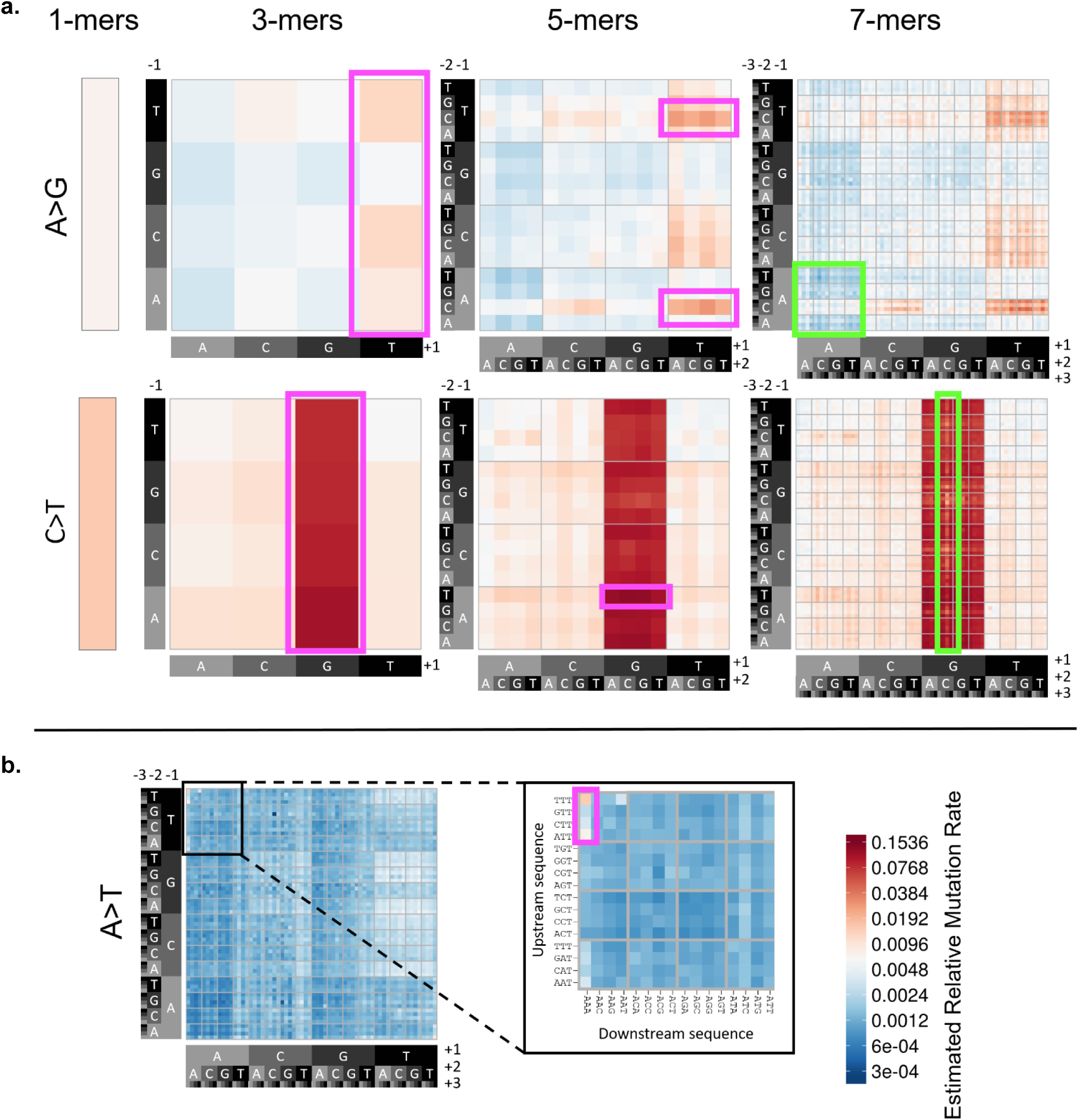
**(a)** Heatmap of estimated relative mutation rates for all possible for A>G and C>T transition subtypes, up to a 7-mer resolution (High-resolution heatmaps for all possible subtypes are included in **Supplementary Fig. 1**). The leftmost panels show the relative mutation rates for the 1-mer types, and the subsequent panels to the right show these rates stratified by increasingly broader sequence context. Each 4x4 grid delineates a set of 16 subtypes, defined by the upstream sequence (y-axis) and downstream sequence (x-axis) from the central (mutated) nucleotide. Boxed regions indicate motifs previously identified by Aggarwala and Voight as hypermutable (pink) or hypomutable (green), relative to their similar subtypes. **(b)** Zoomed-in view showing hypermutable NTT[A>T]AAA subtypes relative to other 7-mer A>T subtypes.

Overall, the 7-mer relative mutation rates estimated from the full set of BRIDGES ERVs span a >400-fold range from 0.0003 (CGT[A>T]CCG) to 0.1416 (ATA[C>T]GCA). For each of the 96 3-mer subtypes, we found overwhelming evidence for heterogeneity in the relative mutation rates among their 16 respective 5-mer constituents (chi-squared tests; all *P* < 10^−231^). Further, 1522 (99%) of the 1536 5-mer subtypes had significantly heterogeneous rates among their respective 7-mer constituents (chi-squared tests; *P* < 0.05) (**Methods**).

### Mutation signatures differ between ERVs and common polymorphisms

We next compared the 7-mer relative mutation rates, estimated either from the BRIDGES ERVs or 1000G intergenic SNVs, to determine if the previously reported patterns of context-dependent mutation rate heterogeneity were consistent with trends observed using ERVs. Across all 24,576 7-mer mutation types, relative mutation rates were highly correlated between the two sets of estimates (Spearman’s r=0.95; **Fig. 2a**). However, when stratified by mutation type, these correlations were often much weaker (r=0.42 to 0.92; **Fig. 2b**). At the individual 7-mer subtype level, discrepancies between the estimated rates were even more pronounced, with 13% of 7-mer subtypes showing differences of 50% or more between the two estimates after normalization. This discordance did not appear to occur randomly across subtypes, as we would expect if these differences were purely stochastic. Instead, we found that subtypes that shared similar flanking sequences often exhibited common patterns of dissimilarity in the estimated rates (**Fig. 2c**). For example, relative mutation rates for C>A and C>G transversions at CpG dinucleotides were respectively 26% and 39% higher in the 1000G estimates compared to the ERV-derived estimates (**Fig. 2c; Supplementary Fig. 2**). Differences in relative mutation rate estimates for A>C and A>G subtypes were also affected by sequence context: we found that the 1000G-derived estimates tended to be significantly higher than ERV-derived estimates among high-GC motifs (4-6 G/C bases in the +/-3bp flanking sequence) compared to low-GC motifs (3 or fewer flanking G/C bases) (t-tests; *P* < 8.0 × 10^−30^) (**Supplementary Fig. 2**; **Supplementary Table 3**). This observation is consistent with the known correlation between GC content and biased gene conversion^18, 31^, though other evolutionary processes may also have contributed.

**Figure 2.**
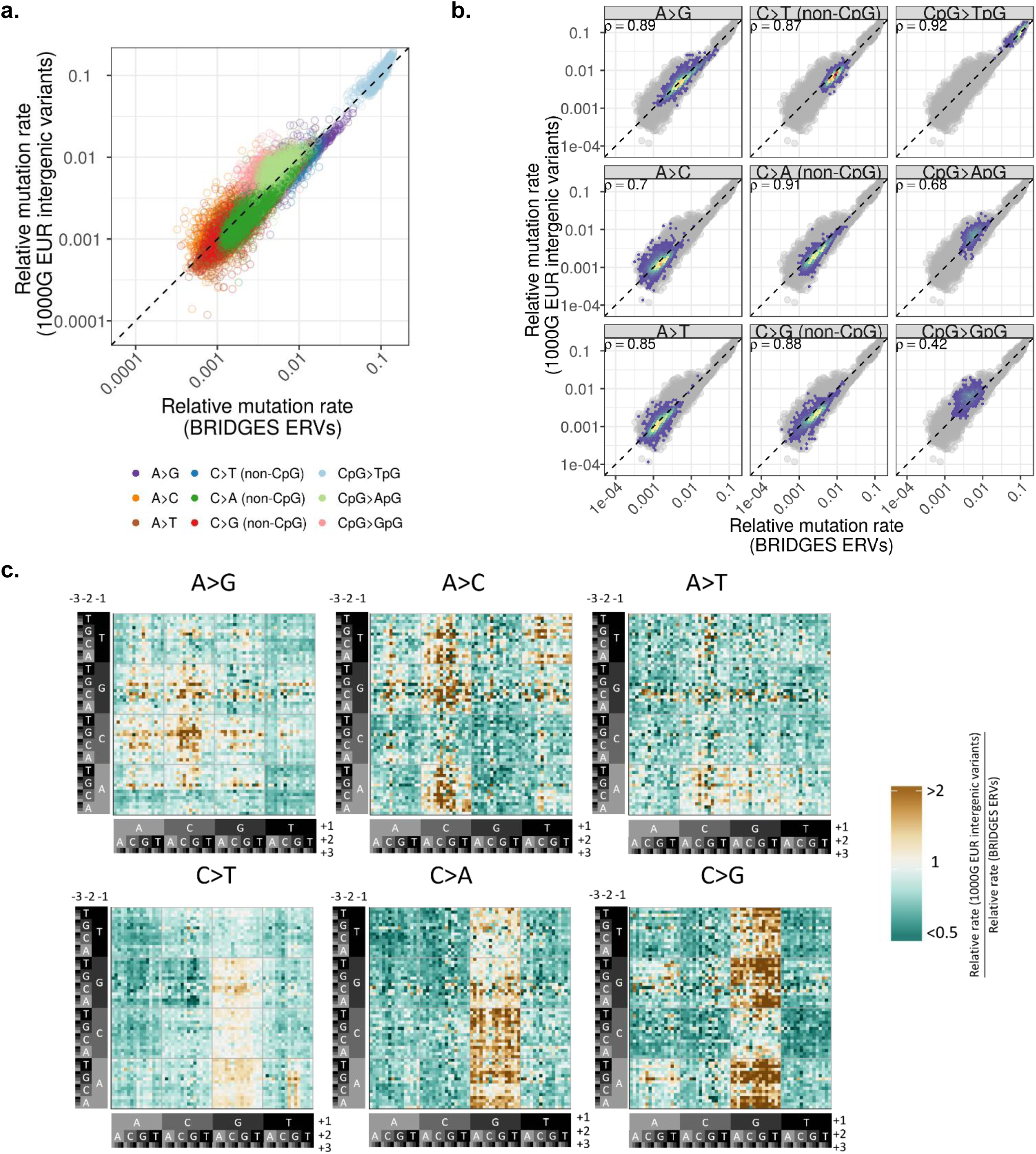
**(a)** Relationship between 7-mer relative mutation rates estimated among BRIDGES ERVs (x-axis) and the 1000G intergenic SNVs (y-axis) on a log-log scale. We note that the strength of this correlation is driven by hypermutable CpG>TpG transitions. **(b)** Type-specific 2D-density plots, as situated in the scatterplot of **(a)**. The dashed line indicates the expected relationship if no bias is present. **(c)** Heatmap showing ratio between the relative mutation rates for each 7-mer mutation subtype. Subtypes with higher rates among the 1000G SNVs (relative to ERV-derived rates) are shaded gold, and subtypes with lower rates in the 1000G SNVs are shaded green. Relative differences are truncated at 2 and 0.5, as only 2.5% of subtypes showed differences beyond this range.

We considered the possibility that these patterns of dissimilarity were simply due to technical differences between the BRIDGES and 1000G samples (e.g., sequencing platform, variant calling and QC methods, sample demography). To address this concern, we estimated relative mutation rates using 12,088,037 variants with a minor allele count ≥10 (MAC10+) in the BRIDGES sample and compared these estimates to the ERV-derived and 1000G-derived estimates (**Supplementary Note**). Importantly, the MAC10+ and 1000G-derived relative mutation rate estimates were more similar to each other both across all types combined (r=0.98; **Supplementary Fig. 3a**) and within each mutation type (r=0.87-0.98; **Supplementary Fig. 3b**), whereas differences between the MAC10+ and ERV-derived estimates agreed with what we observed between the 1000G and ERV-derived estimates (overall: r=0.95; **Supplementary Fig. 4a**; type-specific: r=0.45-0.95; **Supplementary Fig. 4b**). We also found the same sequence-specific patterns of discordance between the ERV and MAC10+ estimates as we did when comparing the ERV and 1000G estimates, with MAC10+ data showing higher rates of CpG transversions and A>G/A>C mutations in GC-rich motifs (**Supplementary Fig. 4c**), but between the MAC10+ and 1000G estimates, these differences were absent or much weaker (**Supplementary Fig. 3c**).

Collectively, these results suggest that the dissimilarities between ERV-based and common SNV-based relative mutation rate estimates are driven not by differences in the data source or analysis pipeline, but by differences in the allele frequencies of the variants used to estimate the rates. There are two plausible explanations for these differences: either 1) the ancestrally older variants included in the 1000G data are under the influence of evolutionary processes that have altered the relative frequencies among subtypes, or 2) even after our careful data cleaning and filtering, certain sequence motifs are enriched for false positive or false negative sequencing errors in the BRIDGES ERVs.

These two scenarios can be tested by comparing how well each set of relative mutation rate estimates describes the observed distribution of true *de novo* mutations. We reasoned that if biased sequencing errors have occurred, such spurious effects would occur more frequently among BRIDGES ERVs, as errors would need to be present in multiple individuals to manifest among the common variants included in the 1000G data. In such a scenario, we would expect the 1000G estimates to explain the distribution of true *de novo* mutations more accurately. In contrast, if the relative mutation rate estimates have been influenced by evolutionary processes, such biases should have a stronger effect on the 1000G estimates and the ERV-derived estimates would provide a better fit.

### Distribution of *de novo* mutations is predicted more accurately by ERVs than common variants

We implemented this validation strategy by comparing how accurately different sets of relative mutation rate estimates predicted the incidence of 46,813 bona fide *de novo* mutations collected from two family-based WGS datasets: The Genomes of the Netherlands (GoNL) project^9^ and the Inova Translational Medicine Institute Preterm Birth Study^12^ (ITMI) (**Methods; Supplementary Fig. 5**). We set these *de novo* mutations against a randomly-selected background of 1 million non-mutated sites, then applied logistic regression models where we used each set of relative mutation rate estimates (either ERV-based estimates at varying K-mer lengths, or 1000G-based 7-mer estimates) to predict the log-odds of observing a *de novo* mutation at each of the 1,046,813 sites. We evaluated the performance of each model by calculating two likelihood-based goodness-of-fit statistics: the Akaike information criterion (AIC), and Nagelkerke’s pseudo-R^2^ (**Methods**).

We first compared the AIC of prediction models based on either the 1-mer, 3-mer, 5-mer, or 7-mer ERV-based relative mutation rate estimates to confirm whether broader motifs truly improve the ability to predict *de novo* mutations. As shown in **Table 1**, goodness-of-fit improved consistently with consideration for longer motifs, with the ERV-based 7-mer model producing the best fit overall. To assess if our results are affected by mapping artifacts, we also re-estimated the ERV-based 7-mer relative mutation rates after applying the 1000 Genomes strict accessibility mask (**Supplementary Note**). We note that the masked and unmasked 7-mer rates are highly concordant, and most discrepancies appear to be an artifact of sampling variation due to fewer ERVs in the masked data (**Supplementary Fig. 6**). When applied to predict the *de novo* mutations, these masked rates decreased model performance slightly compared to the unmasked 7-mer model (**Table 1**), suggesting that reducing the number of observed ERVs has a larger effect on the precision of our estimates than any motif-specific calling error in hard-to-map regions of the genome. These trends did not change when using fewer or more non-mutated sites (**Supplementary Table 4**) nor when applied exclusively to either the GoNL or ITMI mutations (**Supplementary Table 5**), indicating the regression was not merely fitting to cryptic errors in the validation data. We next analyzed each mutation type separately to determine if the same trend of improved goodness-of-fit using longer K-mers was true for different mutation types. In each of these type-specific validation models, the ERV-based 7-mer relative mutation rate estimates provided a significantly better fit than estimates in smaller K-mers (**Supplementary Table 6**).

**Table 1.** Goodness-of-fit statistics for mutation rate estimates applied to*de novo* testing data.

| Mutation rate estimation strategy | | | AIC | $\Delta AIC^\dagger$ | AIC rank* | Nagelkerke's $R^2$ |
| --- | --- | --- | --- | --- | --- | --- |
| Subtype length | Study | Variant type |  |  |  |  |
| 1-mers | BRIDGES | ERVs | 353,896 | 0 | 7 | 0.088 |
| 3-mers | BRIDGES | ERVs | 343,716 | -10,180 | 5 | 0.118 |
| 5-mers | BRIDGES | ERVs | 341,778 | -12,118 | 3 | 0.124 |
| 7-mers | BRIDGES | ERVs | 341,295 | -12,601 | 1 | 0.126 |
| 7-mers | BRIDGES | ERVs<br>(passing 1000G strict mask) | 341,484 | -12,412 | 2 | 0.125 |
| 7-mers | BRIDGES | MAC10+ | 342,886 | -11,010 | 4 | 0.121 |
| 7-mers | 1000G | Intergenic SNVs<sup>7</sup> | 344,003 | -9,893 | 6 | 0.118 |

<sup>†</sup>difference in AIC from the baseline BRIDGES 1-mer model
\*lower AIC rank indicates better model performance

We then compared the goodness-of-fit of logistic regression models using either BRIDGES ERV-based or 1000G intergenic SNV-based 7-mer relative mutation rate estimates. Across all types combined, the 1000G 7-mer model predicted the *de novo* mutations less accurately than all ERV-based models except the baseline 1-mer model (**Table 1**). Considering different mutation types (**Supplementary Table 6**), we observe that for A>C and A>G mutations, the 1000G 7-mer rates provide a worse fit than ERV-derived 5-mer rates; for A>T mutations the 1000G fit is even worse than ERV-derived 3-mer rates. For all C>N mutations except CpG>GpG transversions, the 1000G rates provides a better fit than ERV-derived 5-mer rates; for CpG>GpG mutations the 1000G rates again fit slightly worse than ERV-derived 3-mer rates. These results thus support a scenario in which ancestrally older variants have been influenced by evolutionary biases, and do not reflect patterns of mutation rate heterogeneity observed among true *de novo* mutations as accurately as ERVs.

### Effects of genomic features vary by mutation type and sequence context

Family-based sequencing studies have been instrumental in identifying genomic features that are associated with variation in the germline mutation rate^9,11,25^. However, these studies have only described the marginal effects of features on the entire spectrum of mutation, and have not assessed if the effect of a genomic feature might vary according to the local sequence context. To determine how the distribution of recent mutations varies with respect to the genomic landscape, we selected 14 genomic features (**Supplementary Table 7**) and estimated the joint effects of these features on the mutation rate of each 7-mer subtype using multiple logistic regression, where the dependent variable is the presence or absence of an ERV centered at a given sequence motif (**Methods**). Subtypes with few observed ERVs have little power to detect significant associations, so we estimated the effects of features only for the 24,396 of 24,576 (99.3%) 7-mer subtypes with at least 20 observed ERVs, resulting in 392,128 parameter estimates (**Supplementary Table 8; Supplementary Fig. 7**). We note that >84% of the 7-mer subtypes we evaluated contained >10 times as many ERVs as parameters estimated, so these estimates are unlikely to be an artifact of overfitting. To identify significant effects among the many associations tested, we applied a false discovery rate (FDR) cutoff of 0.05 to the p-values for each feature across all subtype-specific estimates. Of the 24,396 7-mer subtypes analyzed, 3,481 had at least one genomic feature significantly associated with mutability, with 6,152 significant associations among the 392,128 tests.

Three features (H3K9me3 peaks, recombination rate, later replication timing) were associated with higher relative mutation rates across nearly all significantly associated 7-mer subtypes (**Fig. 3a**), consistent with previously reported mutagenic effects of these features: cancer studies have shown that H3K9me3 marks are one of the strongest predictors of somatic SNV density^32, 33^, and recombination and late replication timing are both known to correlate with increased germline mutation rates^8, 9^. In addition, four features (H3K36me3 peaks, DNase hypersensitive sites [DHS], GC content, CpG islands) were each associated with both higher and lower relative mutation rates, depending on the mutation type and, in some cases, the sequence motif. These features have been previously implicated in variation in germline or somatic mutation rates, but only as marginal effects, not type- or subtype-specific. H3K36me3 has been shown to regulate DNA mismatch repair machinery *in vivo*^34^. DNase hypersensitivity was previously reported to be associated with increased germline mutation rates^25^,though cancer genome studies have claimed DHS are susceptible to both increased and decreased somatic mutation rates^35, 36^. CpG islands were associated with ∼3-fold lower mutation rates in 99% (1015/1024) of CpG>TpG 7-mer subtypes, consistent with known patterns of DNA hypomethylation in CpG islands^37^, but are associated with higher relative mutation rates in subtypes of other types.

**Figure 3.**
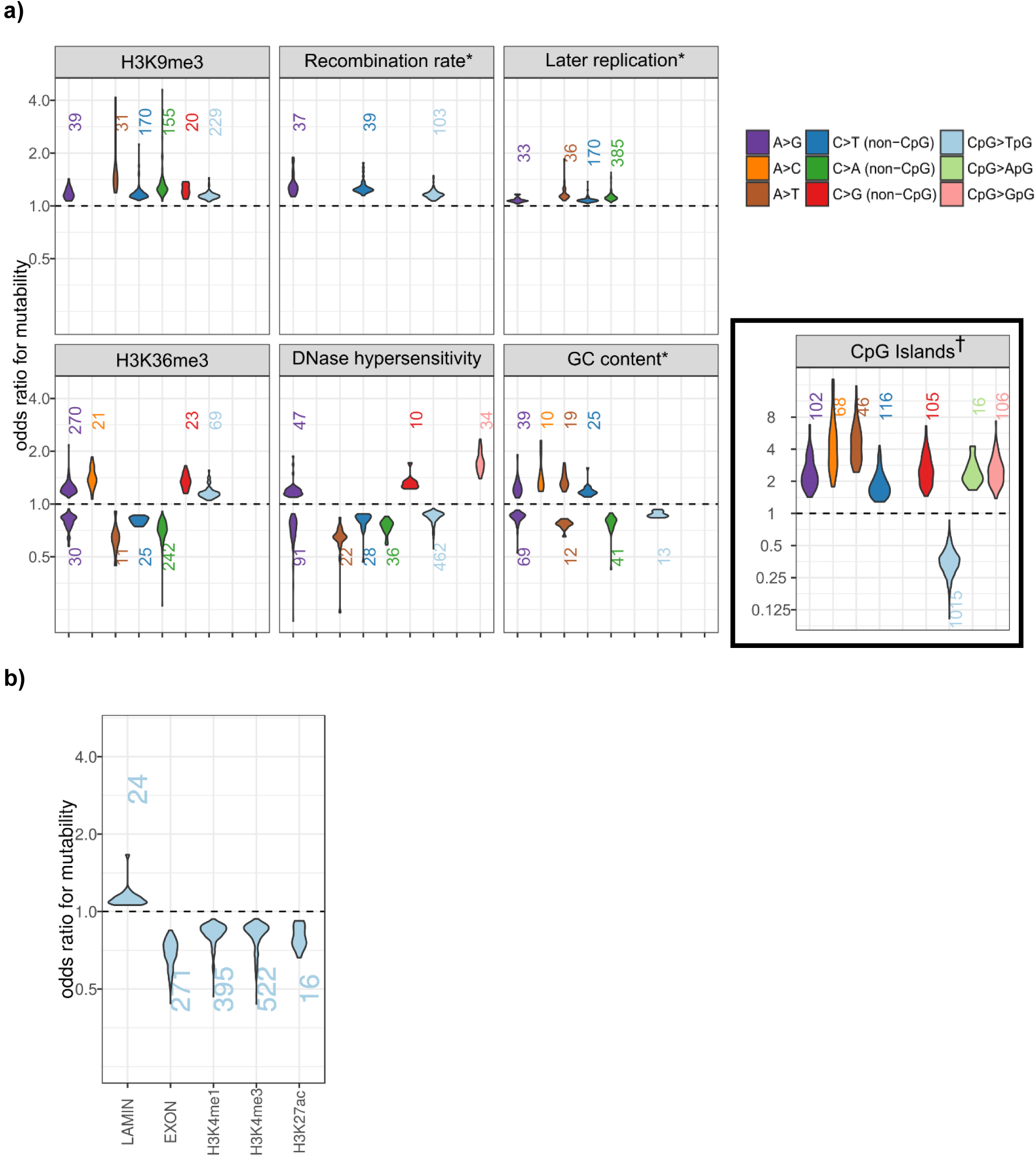
**(a)** Distributions of statistically significant mutagenic effects for 7 genomic features where associations with multiple mutation types were detected. For features with bidirectional effects, we separately plotted distributions of positive associations (OR > 1; above dashed line) and negative associations (OR < 1; below dashed line). The number of 7-mer subtypes within each type for which that feature is statistically significant in a positive or negative direction is shown above or below each distribution. Distributions are only shown for types with 10 or more 7-mer subtypes associated in the same direction. *Odds ratios for the 3 continuously-valued features (recombination rate, replication timing, and GC content) indicate the change in odds of mutability per 10% increase in the value of that feature. ^†^Effects in CpG islands are tend to be stronger than other features, so are shown on a wider scale. **(b)** Distributions of significant mutagenic effects for the 5 features only associated with CpG>TpG transitions.

Finally, for CpG>TpG transition subtypes, lamin-associated domains were associated with higher relative mutation rate and three histone marks (H3K4me1, H3K4me3, and H3K27ac) were associated with lower relative mutation rates (**Fig. 3b**). These results are consistent with published findings of correlations between these features and DNA methylation: lamin-associated domains were previously found to associate with focal DNA hypermethylation in colorectal cancer^38^, and H3K4me1, H3K4me3, and H3K27ac are known markers of DNA hypomethylation^39–41^. We also found that exons were associated with lower relative mutation rates for several CpG>TpG subtypes (**Fig. 3b**), which is in line with findings of lower somatic SNV density in gene-rich regions^32^, though it is unclear if this is also driven by DNA hypomethylation.

### Estimated effects of local genomic features predict *de novo* mutations

We applied these 7-mer+features mutation rate estimates to predict the set of GoNL/ITMI *de novo* mutations, using the same evaluation framework by which we compared the performance of the estimation strategies we described earlier. Model fit statistics indicated that the estimates based on both 7-mer sequence context and genomic features describe the distribution of *de novo* mutations significantly better than the 7-mer-only estimates (**Fig. 4**). When partitioned by mutation type, we find that inclusion of genomic features improves model fit for 8 of the 9 basic mutation types. These differences tend to be weaker among transversion types, likely because there were fewer *de novo* mutations of these types available (**Fig. 4; Supplementary Table 6**). Including genomic features had the largest effect on the prediction of CpG>TpG transitions, consistent with the expected associations between certain features and DNA methylation.

**Figure 4.**
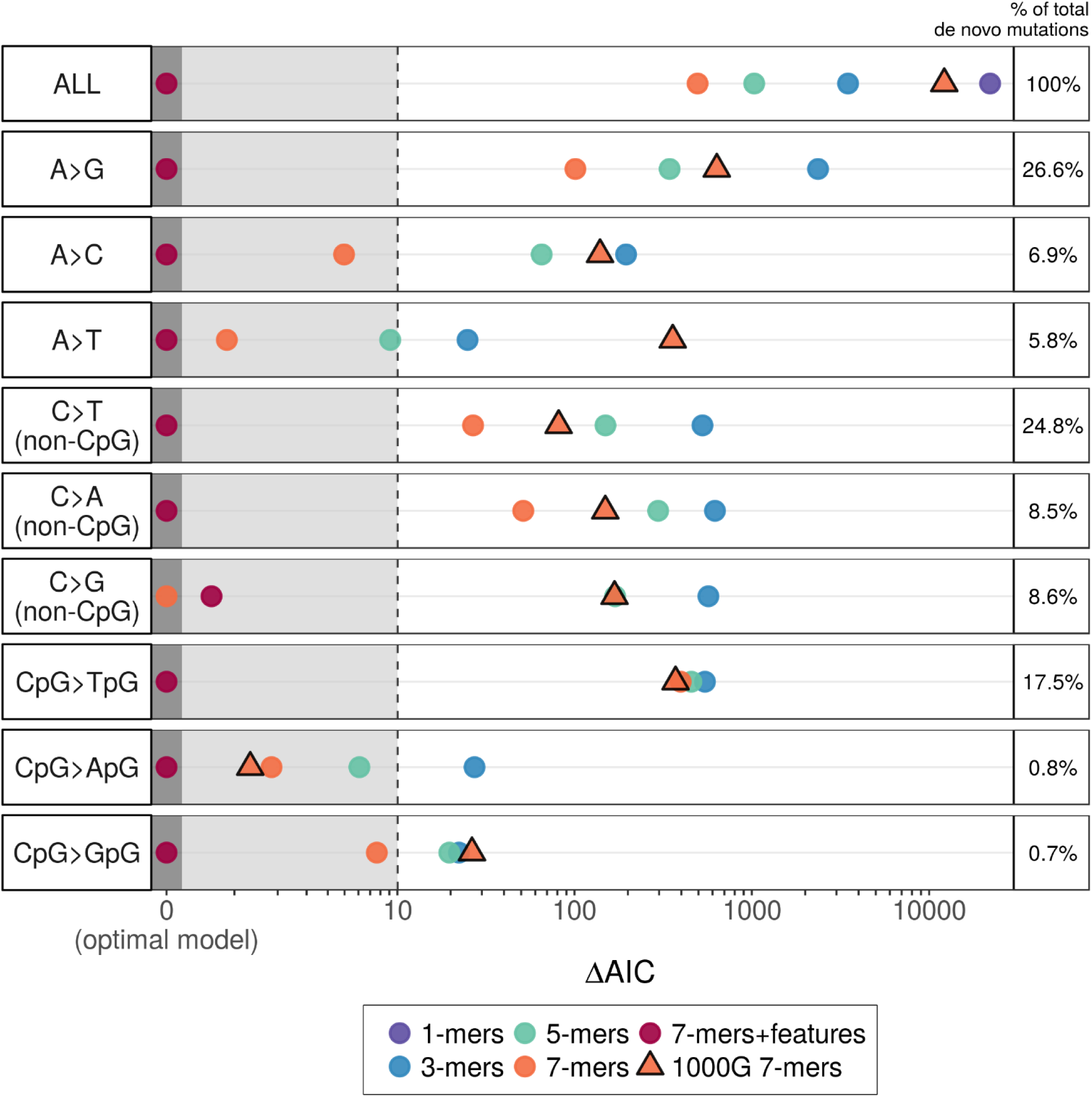
Comparison of goodness-of-fit for different mutation rate estimation strategies, applied to predict the GoNL/ITMI *de novo* mutation data. For each mutation type and each model *i*, we calculated Δ*AIC*_*i*_ = *AIC*_*i*_ - AIC*_min_* as a measure of relative model performance, with lower values of Δ*AIC* indicating better fit. Δ*AIC* is shown on the horizontal axis on an arcsinh scale. For each mutation type, the best-fitting model thus has a Δ*AIC* = 0. Models with Δ*AIC* < 10 (grey-shaded area) are considered comparable to the optimal model, whereas models with Δ*AIC* > 10 are considered to explain substantially less variation than the optimal model^42^.

We also looked to verify that the subtype-specific effects of genomic features, as estimated using the BRIDGES ERVs, were also observed in actual *de novo* mutations. For each of the features, we identified all GoNL/ITMI *de novo* mutations occurring in the set of 7-mer subtypes found to be significantly associated with that feature. We then tested if the subtypes associated with a given feature contained an enrichment or depletion of *de novo* mutations inside versus outside of regions covered by that feature (**Methods**). If a feature was found to have positive effects for certain subtypes and negative effects for others, we separated subtypes by the direction of effect. As shown in **Supplementary Table 9**, 10 of the 20 tests were statistically significant in the expected direction (chi-squared tests; *P* < 0.05), confirming that many of the subtype-specific effects estimated using ERVs are operative among true *de novo* mutations.

### Subtype-specific effects reveal potential mechanisms of hypermutability

The fine scale variability of mutation rate captured by our approach can potentially indicate granular context-dependent mutation mechanisms in the germline. Here we describe two examples. Recent studies of various cancers revealed an elevated somatic mutation rate in transcription factor binding sites within DNase hypersensitive sites (DHS), likely caused by inhibition of nucleotide excision repair machinery^36,43,44^. One of the most common transcription factor binding sites in the genome is the 5’-CCAAT-3’ motif, which is targeted by a family of transcription factors known as CCAAT/Enhancer Binding Proteins (CEBPs)^45^. Because CEBP binding sites were found to be significantly enriched for somatic mutations in multiple cancer types^43^, we speculated that a similar mechanism may be operative in the germline. Adjusting for other genomic features, our analysis indeed shows DHS are significantly enriched for A>G (but not A>C or A>T) ERVs at four of the 16 possible CCA<u>A</u>TNN motifs (1.1 to 1.3-fold enrichment; Wald test; *P* < 2 × 10^−4^). Consistent with the significant effect detected using ERVs, we found that the rate of CCA[A>G]TNN *de novo* mutations in the GoNL/ITMI dataset was 1.7-fold higher when occurring within DHS versus non-DHS regions (1-df chi-squared test; *P* < 0.0055).

A second example are the previously mentioned 5’-NTT<u>A</u>AAA-3’ motifs, which harbor A>T mutations at a ratê6.1-fold higher than the background (1-mer) A>T rate (**Supplementary Table 2d**). However, in ATT<u>A</u>AAA or TTT<u>A</u>AAA motifs occurring in DNase hypersensitive sites, the mutation rate is reduced by over 3-fold (Wald test; *P* < 2.8 × 10^−22^). The TTAAAA hexamer is the primary insertion target for LINE-1 retrotransposons and *Alu* elements^46^, and is known to be nicked by L1 endonuclease (L1 EN) at the TpA dinucleotide, even when no retrotransposition takes place^47^. Moreover, the rate of L1 EN-induced damage has been shown to vary according to the nucleosomal context of target motifs^48^, consistent with our finding that the NTT[A>T]AAA mutation rate differs in DHS. Overall, this pattern of sequence- and feature-dependent mutability suggests that L1 EN nicks are mutagenic, resulting in A>T transversions. A more detailed analysis of the potential sources behind this mutation signature is presented in the **Supplementary Note**.

## Discussion

The main motivation of our study is to understand the genome-wide variation of germline mutation rates in humans. We bring to this task two innovations: first, we take advantage of large-scale WGS data, focusing on extremely rare variants as a potentially more powerful data source than currently available collections of *de novo* mutations^9,10,12,25^ or common variants^7, 13^. Second, building upon previous attempts to holistically model the relationship between sequence context, genomic features, and mutation rate, we estimate fine-scale mutagenic effects of multiple genomic features. Unlike previous studies, which estimated the impact of genomic features by treating all single-nucleotide mutation subtypes in aggregate^25^, we allow for the possibility that mutation rates of sequence motifs are differentially affected by these features.

Our results not only confirm the previously reported hypermutable effects of specific sequence contexts (e.g., higher A>T mutation rates at NTTAAAA motifs) and genomic features (e.g., higher mutation rates in late-replicating regions^9^), but also demonstrate that feature-associated effects previously only described in somatic cells are also present in the germline (e.g., a positive association with H3K9me3 peaks^32^). Unexpectedly, our approach identifies certain genomic features, such as H3K36me3 peaks, DNase hypersensitive sites, and CpG islands, that may act to both suppress and promote mutability depending on the type of mutation and local sequence context (**Fig. 3**), providing more detailed insight into how the mutation rate is modulated across the genomic landscape.

We note that power to detect a given level of mutagenic effects of genomic features depends on the number of ERVs of a given 7-mer subtype: of the 6,514 significant associations, 93% were detected in 7-mer subtypes with more than 731 ERVs, which is the median number of ERVs among all 7-mer subtypes. Thus, a larger dataset of ERVs will likely reveal even more cases of association, and will enable the study of mutagenic effects within longer sequence motifs, additional genomic features, and interactions or nonlinear effects of these features. Although there is strong theoretical and empirical evidence that the distribution of ERVs is largely unaffected by natural selection^23, 24^, we acknowledge that very strong purifying selection may have reduced the number of ERVs in highly conserved functional regions, so we may have underestimated mutation rates for these loci. We also note several of the genomic features used in our study were assayed in somatic cell lines or aggregated over multiple cell types (**Supplementary Table 7**). The currently available data for these features thus provides only a crude approximation of the true genomic variation in germ cells, so the effects we estimated have likely regressed towards the mean. Generating precise maps of genomic features within germ cells (and throughout the stages of gametogenesis) will be necessary to fully describe how germline mutation rates are influenced by the genomic landscape. Despite these limitations, the context-specific mutation rates and context-feature interactions reported here provide the most accurate map to date of germline mutation variation, as demonstrated by their improved ability to predict genuine *de novo* mutation patterns.

Even without accounting for the effects of genomic features, our ERV-derived mutation rate estimates for 7-mer subtypes are consistently more accurate than those based on mostly common SNVs from 1000 Genomes Project data^7^. Remarkably, even coarser estimates—the ERV-derived 5-mer and 3-mer rates—predict the spectrum of *de novo* mutations more accurately than the 1000G 7-mer estimates, demonstrating the merit of ERVs as a refined data resource for studying innate mutation patterns. This result has two important implications. First, it suggests that many high-frequency variants in presumably neutral regions of the genome likely have experienced biased evolutionary processes, such as selection and gBGC, or these variants may have arisen by past mutational processes that have shifted over time or are no longer active^22^. In either case, we demonstrated that the distribution of ERVs provides a more accurate appraisal of recent or ongoing mutagenic processes than common SNVs. Second, this reaffirms the high quality of ERVs in our data: the potential errors due to calling or mapping biases among these ERVs are likely weaker than the evolution-driven biases affecting the older variants.

Because the germline mutation rate is one of the most critical parameters in the study of genetic variation, we envision a wide range of applications that stand to benefit from incorporating our genome-wide map of mutation rate estimates. Currently, many methods that rely on simulating “baseline” mutations, such as the pathogenicity scoring algorithm *CADD*^49^ and coalescent simulator *ms*^50^, do not account for context-dependent mutation rate differences. Likewise, clinical applications for differentiating disease-causing mutations from background variation require a precise estimate of the expected *de novo* mutation rate, but even the most advanced of these only consider differences in 3-mer or 7-mer sequence contexts, and are based on intergenic SNVs from 1000 Genomes data^7, 51^. Incorporating more accurate sequence- and feature-dependent estimates of mutation rates will likely lead to more realistic simulations and greater confidence in the inferences made by these methods. Another particularly relevant area of research where our results might be applicable is the study of how germline mutation mechanisms have evolved over time^22,52,53^. If mutator phenotypes have frequently come and gone throughout the evolutionary history of humans (as hypothesized by Harris and Pritchard^22^), it seems likely that the effects of mutational modifiers have been extremely subtle, manifesting as granular context-specific mutation signatures. Our results, which describe the present-day pattern of mutation rate heterogeneity in Europeans, thus provide a wealth of potential hypotheses for investigating how these mutation processes have been shaped via past evolution.

To facilitate the use of our genome-wide mutation rate estimates in other analysis and simulation pipelines, we have used our full model to predict the mutation rate at every location in the genome, and created a genome browser track to visualize the predicted mutation rates alongside other genomic data. Ultimately, the refined mutation patterns from ERVs and the detailed dissection of context-feature effects serves as a quantitative foundation for better understanding the molecular origins of mutation rate heterogeneity and its consequences in heritable diseases and human evolution.

## Acknowledgements

Funding for this research was provided by US National Institutes of Health (NIH) grant R01HG005855 (S.Z. and J.L.). J.C. was supported by the NIH/National Human Genome Research Institute Genome Science Training Program (T32HG00040). The BRIDGES study was supported by NIH grants R01MH094145 (M.B., R.M.M.) and U01MH105653 (M.B.). Additional acknowledgements from collaborating members of the BRIDGES consortium are detailed in the **Supplementary Information**.

## Author contributions

J.C., S.Z., J.L., and L.S. wrote the manuscript. J.C., S.Z., and J.L. designed the mutation models. J.C. performed the analyses and created the online annotation utility and interactive heatmap. M. B. and H.M.K. provided critical feedback and evaluation of the manuscript. A.L., M.F., and H.M.K. performed variant calling and filtering of the BRIDGES samples and curated the raw data. The BRIDGES study was designed by A.L., L.S., R.M., and M.B., with sequencing led by S.L. and R.M.

## Methods

### Sample description

The BRIDGES sample contains 3,927 unrelated European American bipolar disorder cases and controls. The cases and controls from the Centre for Addiction and Mental Health (CAMH) in Toronto (n=830), the Institute of Psychiatry, Psychology and Neuroscience (IoPPN) and King’s College London in London, U.K. (n=845)^54^, the Genomic Psychiatry Cohort (GPC) (n=1,151)^55^, and the Prechter Repository (n=363)^56^ were collected as previously described, as were the STEP-BD cases (n=304), obtained from the NIMH repository^57^, and the Minnesota Center for Twin and Family Research (MCTFR) study controls (n=434)^58^. In all studies, DNA was extracted from blood-based samples. All human research was approved by the relevant institutional review boards and conducted according to the Declaration of Helsinki. All participants provided written informed consent.

### Sample library preparation

The concentration of each DNA sample was measured by fluorometric means (PicoGreen, Thermo Fisher, Woburn, MA, USA) followed by agarose gel electrophoresis to verify the integrity of DNA. Six-hundred nanograms of DNA was sheared with acoustic shearing (Covaris, Woburn, MA, USA) to an average size of 400nt. Following shearing, the samples are transformed to a sequencing library using standard protocols to create a paired-end library. Briefly, sheared DNA was end-repaired, A-tailed and ligated with Illumina adaptors (New England Biolabs, Ipswitch, MA, USA). Following ligation, indexed primers were used to amplify the final libraries for each sample. Each sample received two indexes: 96 i7 indexes were used to identify each sample in each 96-well reaction plate while a single i5 index was used for each plate. This combination of indexes uniquely coded all samples in the project when both the i7 and i5 indexes were read during sequencing. Following six cycles of PCR (Kapa Biosystems, Wilmington, MA, USA), libraries were purified and quality controlled by assaying the final library size using the Agilent Bioanalyzer (Agilent Technologies, Santa Clara, CA, USA) and quantitating the final library via real-time PCR (Kappa Biosciences). A single peak between 300-400bp indicates a properly constructed and amplified library ready for sequencing. PCR cycles for amplification are kept to a minimum to minimize PCR duplication rate and maximize library complexity.

### Sequencing

Sequencing was performed per Illumina protocol, essentially as described by Bentley et al.^40^. Libraries were pooled in sets of 12 samples and each pool sequenced on a single lane of a HiSeq 2500 flowcell using version 3 Illumina chemistry at paired-end 100nt read lengths. Each library pool was loaded at 13pM to generate 160-180M paired reads per lane. Multiple flowcells of the library pools were performed to generate a final data set with an average coverage of 9.6x per sample.

### Sample filtering and data quality control

Among the 3,927 samples attempted, three failed library preparation and were not sequenced. We removed an additional 162 samples due to quality issues: five with imbalanced read counts between read 1 and read 2, four with improperly generated BAM files, 16 that had an average coverage <3x, and 137 due to high contamination (FREEMIX or CHIPMIX score >3% using VerifyBAMID^59^). For samples that failed for multiple reasons, we report a single category for simplicity.

Among these 3,762 samples, reads were mapped to Build 37 of the human reference genome (including decoy sequence^28^), with alignment and variant calling performed using the GotCloud pipeline^60^. After variant calling, we applied additional sample-level filtering as described below to obtain the 3,716 included in our analysis. We first excluded 10 case samples that were not phenotyped as type 1 bipolar disorder (removed solely for consistency with ongoing analyses of the BRIDGES data that do require phenotypes). We identified and removed an additional 23 samples that showed evidence of sample swaps in VerifyBAMID^59^, but had not been excluded from variant calling. We next computed continental-ancestry PCA coordinates by projecting BRIDGES samples in the coordinate space of the 1000 Genomes phase 1 samples^61^. We dropped 11 samples identified as PC ancestry outliers, defined by PC1<0.01 or PC2<0.025. We then checked for relatedness using the 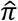 statistic (i.e., estimation of pairwise identity-by-descent based on LD-pruned SNPs), computed in plink^62^. Nearly all pairwise sample comparisons were consistent with being unrelated, with 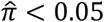 for 99.9% of sample pairs. Two samples were dropped due to relatedness, as the 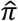 between these was 0.5, indicating the two were full siblings.

These filters reduced the sample to 3,716 individuals, in which we called 37,470,516 autosomal singleton SNVs in the mappable genome (i.e., non-N reference bases in the GRCh37 reference genome) that passed the variant-level filtering criteria implemented in the GotCloud pipeline^60^. Prior to performing our analyses, we examined how these 37.5 million ERVs were distributed across individual samples to identify and remove individuals that showed abnormal patterns of variation due to systematic sequencing errors or batch effects. In brief, we adapted the non-negative matrix factorization (NMF) technique described by Lawrence et al.^63^ to summarize the distribution of ERVs unique to each individual as a composite of 3 distinct “signatures.” For each of the 3,716 individuals in our sample, we calculated a vector of 96 3-mer relative mutation rates (described below) using only the ERVs observed in that individual, generating a 3,716 x 96 rate matrix. Decomposition of this matrix via NMF produces a 3,716 x 3 matrix describing the relative contribution of each signature to the observed mutation spectrum per individual. Because we assume the relative mutation rate of any given subtype should be similar across individuals, it follows that the contribution of a given NMF signature should also be similar. We removed 156 individuals where one or more signatures had a contribution >2 standard deviations away from the mean contribution of that signature calculated across all individuals, reasoning that ERVs observed in these individuals are more likely to be errors. The final sample used in our analyses thus consists of 3,560 individuals, in which we identified 35,574,417 singletons. Additional details of this filtering strategy are described in the **Supplementary Note**.

### Mutation subtypes and calculation of relative mutation rates

Each of the 35,574,417 singletons can be classified into one of 6 basic mutation types, defined by the reference and alternative allele: A>C, A>G, A>T, C>T, C>G, and C>A. The notation of A>C includes both A-to-C mutations and complementary T-to-G mutations. For each mutation type, we further define a set of mutation subtypes by the bases flanking the variant site. Since there are 4 possible bases at both the +1 position and the -1 position, there are 4x4=16 possible 3-mers containing each basic mutation type at the central position, producing 6x16=96 3-mer subtypes. Likewise, there are 6x4^4^=1,536 5-mer subtypes, and 6x4^6^=24,576 7-mer subtypes. To simplify notation, we denote a subtype by the sequence motif containing either an A or a C as the reference base at the central position (e.g., either CGT[A>X]TCG or CGT[C>X]TCG).

For each K-mer subtype, we divided the number of ERVs observed at the central position of the K-mer by the number of times the K-mer is seen in the mappable autosomal regions of the reference genome; we term this proportion the *estimated relative mutation rate*. K-mers in the reference genome were counted by a 1-bp sliding window, so that every possible occurrence of that K-mer was accounted for (e.g., a run of 4 As is counted as two AAA 3-mers shifted by one base). For example, we observed 7,548 C>T or G>A autosomal singletons occurring in an ATA<u>C</u>GCA or TGC<u>G</u>TAT 7-mer motif (the underlined base indicates the variant site) and there are 53,314 such motifs in the autosomal reference genome where this subtype of mutation could be observed, yielding a relative mutation rate estimate of 7,548/53,314=0.1416 for the ATA[C>T]GCA subtype.

### Testing for heterogeneity of relative rates among nested subtypes

As each K-mer can be split into 16 possible (K+2)-mers that share the same internal motif but differ in their terminal bases, the relative mutation rate for each K-mer subtype is the weighted mean of the rates found among its 16 possible (K+2)-mer constituent subtypes. To assess the heterogeneity of relative mutation rates among each set of 16 (K+2)-bp constituent subtypes that share the same K-bp motif, we performed a chi-squared test for uniformity of these rates, with each test having 15 degrees of freedom.

### Mutation prediction model and validation

To evaluate the accuracy of different mutation rate estimation strategies, we applied the estimated rates to predict the incidence of 46,813 *de novo* mutations using logistic regression. These *de novo* mutations were published by two independent studies: 11,020 *de novo* mutations detected in 258 Dutch families by the Genomes of the Netherlands (GoNL) project^9^, and 35,793 *de novo* mutations from 816 families sequenced by the Inova Translational Medicine Institute (ITMI) Premature Birth Study^12^. We combined the observed mutations with 1 million randomly selected sites from the mappable autosomal regions of the reference genome to serve as a non-mutated background, reasoning that ∼20 non-mutated sites for each actual de novo mutation would be sufficient to minimize sampling noise in the set of non-mutated sites; we also repeated this procedure with 500,000, 2 million, and 3 million randomly selected sites to tell if the trends we observed were affected by the size of the non-mutated background. Because each non-mutated site can be ambiguously considered as the background for 3 different mutation types, we divided the 1 million non-mutated sites into 3 non-overlapping sets. We designated A/T and C/G reference bases in the first set (consisting of 333,334 unique sites) as non-mutated A>G and C>T types, respectively, and so on for the second set (A>C or C>G types), and the third set (A>T or C>A types), each of which contained 333,333 unique sites. Hence, we considered a total of 1,046,813 testing sites (1,000,000 unmutated sites and 46,813 de novo mutations), each with one possible mutation event, in our prediction models.

Now let ***i*** = {**1**, …, **1046813**} be an index for the 1,046,813 testing sites. We coded ***d***_***i***_ = **1** if site ***i*** is a de novo mutation and ***d***_***i***_ = **0** otherwise. If a set of estimated relative mutation rates reflects the underlying mutation process, we expect that the odds of a given site for carrying a *de novo* mutation increases with the estimated relative mutation rate of that site. To asses this expectation for all sets of mutation rate estimation strategies (e.g., ERV-based or 1000G-based 7-mer estimates), we annotated each testing site ***i*** with the relative mutation rate estimated under strategy ***M*** (***r***_***i***,***M***_), and used logistic regression to model the probability of a *de novo* mutation at each site as a function of these rate estimates, where ***α***_**0**_ is the intercept term and ***α***_**1**_ is the regression coefficient:

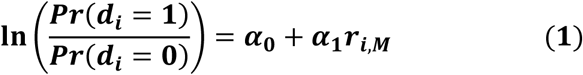

The probability of a mutation at each testing site can then be calculated as:

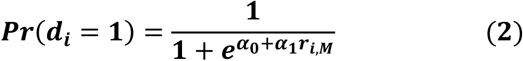

The overall likelihood of model ***M***, given the observed data, is the product of the probability values over all 1,046,813 sites:

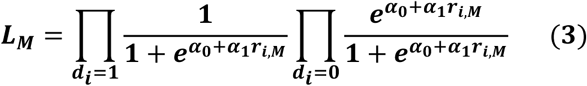

Using this likelihood, we evaluated model fit by the Akaike Information Content (AIC), where ***p*** is the number of parameters in equation (1) (because all models are based on a single covariate of mutation rates, ***p*** = **1** in all cases):

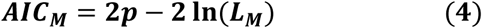

For each model, we also calculate Nagelkerke's R^2^:

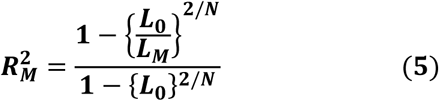

Here, ***L***_**0**_ is the likelihood of a null intercept-only model with no covariates.

Because these likelihood-based goodness-of-fit statistics are calculated across all the basic mutation types combined, they do not provide information about which types benefit most strongly from using expanded sequence motifs. For example, it is possible that any improvement to the overall goodness-of-fit is elicited by context-dependent heterogeneity of a single mutation type, whereas other types might not be significantly affected by using longer sequence motifs, and do not contribute to the improved model fit. To identify these type-specific trends, we stratified our testing data by each of the basic mutation types. To account for the known hypermutability of cytosine at CpG dinculeotides, we separated C>T, C>G, and C>A mutations into CpG and non-CpG types, for a total of 9 basic mutation types. For each type, we repeated the 3-mer, 5-mer, and 7-mer models on only the sites of that type. Within each set of type-specific models, we again compared the goodness-of-fit using AIC and Nagelkerke's R^2^. Note that because the absolute values of AIC and Nagelkerke's R^2^ are a function of the number of data points included in the model, these statistics cannot be directly compared between type-specific models, where the number of data points vary.

### Estimating the effect of local genomic features

We estimated the effect of 14 genomic features (data sources for these features are described in **Supplementary Table 7**) on the relative mutation rate of each 7-mer subtype using the following logistic regression framework. Let ***K*** be the index across all 7-mer subtypes with 20 or more observed singletons (***K*** ∈ {**1**, …, **24396**}). Let ***j***_***K***_ be the index across all sites that are centered at the 7-mer motif that could produce a mutation of subtype ***K***, and let ***Z***_***j_K_***_ = **1** if the site carries a singleton of subtype ***K*** and ***Z***_***j_K_***_ = **0** otherwise. We annotated each site of the considered subtype for 14 genomic features, generating predictors ***F***_***j_K_***,**1**_, …, ***F***_***j_K_***,**14**_. We treated 11 of these features as binary variables (seven histone marks, lamin-associated domains, CpG islands, DNase hypersensitive sites, exons), setting the predictor ***F***_***j_K_***,***g***_ = **1**, ***g*** ∈ {**1**, …, **11**} if the central site of the motif was inside the specified regions and ***F***_***j_K_***,***g***_ = **0** otherwise. For the 3 continuous features (recombination rate, replication timing, surrounding GC content), we set the predictor ***F***_***j_K_***,***g***_, ***g*** ∈ {**12**, **13**, **14**} to the mean value of that feature in a 10kb window centered at the site. Because the inferred effect of some features may be confounded by correlation with read depth and calling rates (e.g., GC content^64^), we included read depth at the central site of the 7-mer as covariate ***F***_***j_K_***,***DP***_. For each 7-mer subtype ***K***, we then evaluated the effect of the genomic predictors on the log odds of mutability for each site ***Z***_***j_K_***_ using the following logistic regression equation:

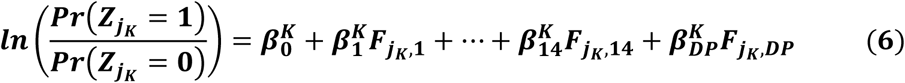

where (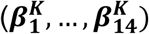) are effects of the 14 considered genomic features on the mutation rate of subtype ***K***, and 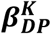 is the effect of the local sequencing depth. The intercept of this model, 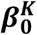, represents the feature-adjusted relative mutation rate for the considered 7-mer subtype. We performed this logistic regression and obtained parameter estimates in R v3.2.3 using the speedglm() function from the *speedglm* package. We performed this procedure for each of the ***K*** ∈ {**1**, …, **24396**} 7-mer subtypes; the resulting beta values and standard errors for 16 x 24,396 estimated parameters are provided in **Supplementary Table 8**. Note that we did not consider estimating interaction effects between the 14 genomic features, as estimating all 2-way interactions would require an additional 14*(13-1)/2=91 parameters per subtype-specific regression, which would lead to overfitting concerns.

To generate a map of mutation rates across the genome, we used the estimated regression coefficients to predict the relative mutation rate (i.e., probability of observing a singleton) at each site ***j*** where a mutation of a given 7-mer subtype could occur:

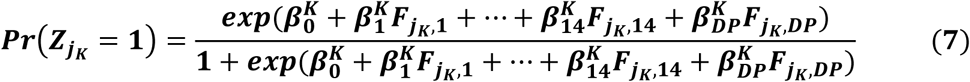

Because there are three possible mutations at every site, we predict 3 independent mutation probabilities (one for each possible alternative allele). For example, for a site centered at a ACGATTG motif, we predict probabilities for A>C, A>G, and A>T alleles, using the parameters estimated from those models. This prediction uses all estimated effects, not just the effects determined to be statistically significant. We note that we did not generate predictions for sites within 5Mb of the start/end of a chromosome, because recombination rate data were not available for these regions^65^.

To assess if inclusion of these genomic features improved upon the 7-mer mutation rate estimates in describing the true distribution of germline mutability, we again tested this model’s ability to predict the known *de novo* mutations from the GoNL^9^ and ITMI^12^ studies. We annotated each of the ***i*** = {**1**, …, **1046813**} testing sites with the predicted mutation rate, ***Pr***(***Z***_***iK***_ = **1**), and calculated the goodness-of-fit using equations 1-5 with this parameter as the predictor. Note that the GoNL/ITMI data included *de novo* mutations within the 5Mb telomeric regions where we could not estimate effects of genomic features. Rather than excluding sites in these regions from our goodness-of-fit comparison, we simply assigned the marginal 7-mer relative mutation rate as the predicted value for these sites, to ensure models were compared using identical data.

### Data availability

We are in the process of submitting the BRIDGES sequence-based genotypes to dbGaP. K-mer-based relative mutation rate estimates are provided in **Supplementary Table 2**. Predicted mutation rates based on sequence context and genomic features at each site have been formatted as a UCSC Genome Browser track, which can be accessed at http://mutation.sph.umich.edu.

### Code availability

All custom scripts used in downstream data processing and analyses are available at https://github.com/carjed/smaug-genetics. A web-based utility and command-line code for annotating a variant call format (VCF) file of genetic variants with estimated 7-mer mutation rates can be accessed at http://www.jedidiahcarlson.com/mr-eel/.

